# Conformational dynamics of ND2 antiporter-like subunit in intact and truncated complex I

**DOI:** 10.64898/2026.10.02.756245

**Authors:** Amiel Abettan, Jonathan Lasham, Erik Endres, Vivek Sharma

## Abstract

Respiratory complex I is a large redox-driven proton pump and contributes to the proton motive force required for ATP generation in mitochondria and bacteria. It has been extensively studied with structural and biochemical approaches leading to enhanced understanding of its three-dimensional architecture. Despite significant progress in deciphering its catalytic function, the redox-coupled proton pumping mechanism remains unclear and extensively debated. The three membrane-bound antiporter-like subunits ND2, ND4 and ND5 are responsible for pumping protons from the negatively charged side (N-side) of the membrane to the positively charged side (P-side). Emerging evidence supports the role of ND5 subunit in releasing pumped protons to the P-side of the membrane, but the question whether ND2 and ND4 also contribute to proton release remains unanswered. Here, we performed atomistic molecular dynamics (MD) simulations of an intact and a truncated model of complex I in which ND5 and ND4 subunits are removed *in-silico*. The simulation data reveals a previously unobserved conformational change in the P-side region of ND2 subunit that occurs much more easily in a truncated model of complex I, as also estimated based on free energy calculations. The conformational change, in part triggered by the artificial exposure of ND2 subunit to lipid bilayer, results in a local hydration of the region, similar to what has been observed in recent cryo-EM structure of plant CI:CIII_2_ supercomplex. We discuss the functional relevance of observed conformational transitions and identify residues that can be manipulated with biochemical approaches to shed light on proton pumping by complex I.

**Highlights:**

* Atomistic MD simulations of intact and truncated respiratory complex I are performed.
* Conformational changes and hydration in membrane-exposed ND2 subunit are observed.
* Functional relevance of conformational changes in proton pumping is discussed.

## 1. Introduction

Respiratory complex I is a large redox-driven proton pump in the respiratory chains of mitochondria and many bacteria [1–3] (Fig. 1A). By catalyzing the reduction of quinone (E_m,7_, midpoint potential at pH 7 ∼ +90 mV) from NADH (E_m,7_ ∼ -320 mV), it converts the redox energy of ∼800 meV to the pumping of up to four protons against an electrical gradient of ∼200 mV across the membrane [4, 5]. In addition to being a key contributor of the proton motive force (ca. 40% of the total) [4, 5], respiratory complex I in mitochondria plays a central role in NAD, quinone (Q) and ROS metabolism [3, 6, 7]. A large number of mitochondrial disease mutations are found within the mitochondrial DNA-encoded proton pumping subunits of complex I [8]. Furthermore, complex I has been implicated in cancer and neurodegenerative diseases [9, 10]. Several small molecule inhibitors that target the Q binding site of complex I have been studied as potential therapeutic agents against neurodegeneration, cancer and other metabolic disorders [11–15]. Due to its physiological importance, complex I has been the focal point of bioenergetics research, and has been explored with biochemical [16–18], biophysical [19–22] and structural approaches [23–26]. Despite extensive investigation, the molecular mechanism of redox-driven proton pumping by complex I remains unknown and extensively debated in the field; see recent reviews [27–30].

**Fig. 1.**
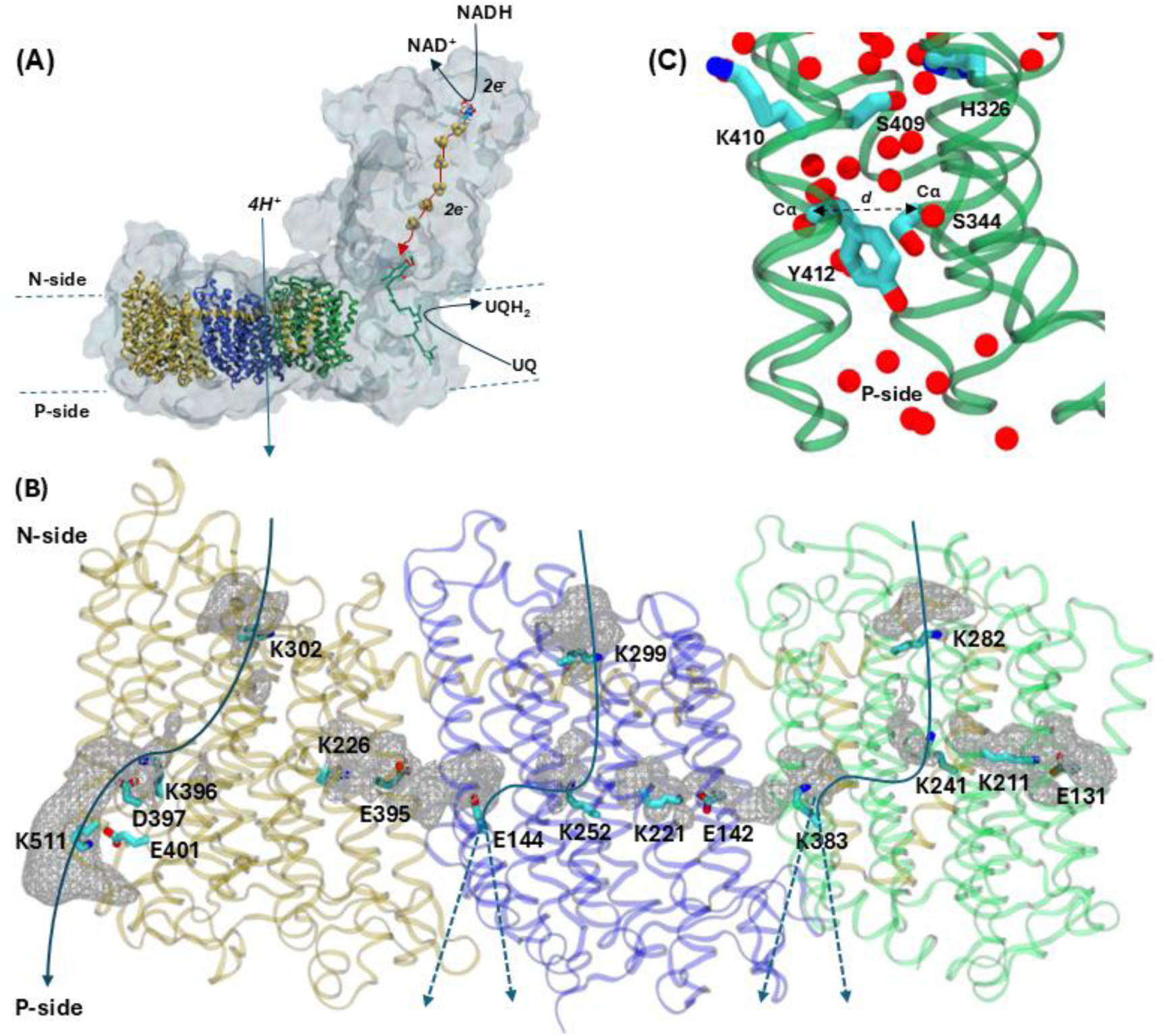
Mitochondrial respiratory complex I. (A) Overall architecture of the mitochondrial complex I from yeast *Yarrowia lipolytica* (PDB 7O71) is shown with core subunits ND2, ND4 and ND5 highlighted in colored ribbon representation. Other subunits are shown with a light grey colored surface. Membrane boundaries are marked with dotted lines. Electron (2 e^-^) and proton (4 H^+^) transfers are shown with red and blue arrows, respectively. (B) Three homologous antiporter-like subunits ND2, ND4 and ND5 are displayed together with the selected titratable amino acid residues and putative proton transfer paths marked. The dotted lines indicate uncertainty in the proton exit routes in ND2 and ND4 subunits for which two distinct routes can be expected; one at the interface of two subunits and another buried within each subunit. Hydration in the three antiporter-like subunits based on 3 μs (3 replicas x 1 μs) of atomistic MD simulation data on high resolution structure of *Y. lipolytica* [29, 31]. The grey mesh represents water occupancy shown at 0.10 isovalue. (C) Water molecules (red spheres) and polar residues closer to the P side of the membrane in ND2 subunit of complex I from *Arabidopsis thaliana* [32] (PDB 8BEF). Selected polar amino acid residues in the region are shown along with the distance *d* marked between Cα atoms of Ser344 and Tyr412 (see main text).

Among the many questions on proton pumping mechanism of complex I [29], one fundamental issue that has remained unresolved is the location of proton pumping routes in its three antiporter-like subunits (ND2, ND4 and ND5, see Fig. 1B). The proton-uptake routes from the N side of the membrane (from cytoplasm or mitochondrial matrix) have been suggested in all three subunits based on high-resolution 3D structures, computer simulations and site-directed mutagenesis studies [27–30]. It is now agreed that the terminal ND5 subunit (Fig. 1B) forms a well-hydrated route that can be used for proton exit towards the P side of the membrane. Whether similar proton release routes exist in the two other subunits ND2 and ND4 remains unclear. It also remains unknown whether the proton exits at the interface of antiporter-like subunits (ND2/ND4 and ND4/ND5 interface) or if the exit route is contained within a single antiporter-like subunit (see Fig. 1, [29]).

Early on, proton pumping function of the antiporter-like subunits of complex I was proposed based on the sequence similarity to Na^+^/H^+^ antiporters [33, 34]. The site-directed mutagenesis data on bacterial complexes [16, 17, 35] provided biochemical support that each antiporter-like subunit participates in proton translocation. The major leap in complex I research occurred when its 3D structures were resolved by X-ray crystallography in a resolution range of 3 to 4 Å. First partial [36, 37] and then a complete structure [38] of bacterial complex I was solved by Sazanov and colleagues. These combined with the structure of mitochondrial complex I from *Yarrowia lipolytica* [39] provided first atomic level insights on the putative proton transfer routes in all three antiporter-like subunits (see Fig. 1B). These structural data agreed well with the earlier mutation analysis of antiporter-like subunits (see above). But due to relatively lower resolution of these structures, water molecules, which are central for proton transfer activity, could not be resolved. MD simulations performed on bacterial complex I structures provided atomistic insights on protein hydration in putative proton transfer pathways of antiporter-like subunits [40, 41]. Already based on these simulation studies, it can be inferred that the proposed proton release routes of ND2 and ND4 subunits (NuoN and NuoM in *E. coli* complex I) are much less hydrated in comparison to ND5 subunit. Later, using the complete structure of *Thermus thermophilus* complex I modeled in a lipid-solvent environment, large-scale atomistic MD simulations were performed [42, 43]. Haapanen and Sharma [42] noted that the proposed proton release route in ND5 subunit was more hydrated than ND2 and ND4. A somewhat similar picture can also be inferred from the data presented by Di Luca and colleagues in ref. [43]. However, transient and weak hydration was observed at the ND2/ND4 and ND4/ND5 interfaces, which was interpreted as potential exit route for protons to the P side of the membrane. A detailed look at the data presented in later simulation studies [44, 45] further corroborates the notion that ND2 and ND4 subunits display weaker levels of hydration than ND5 at the putative proton release sites. These data, now when looked at in hindsight, raises the question whether ND2 and ND4 subunits contain a proton release route or if it is only the ND5 subunit that is responsible for releasing protons to the P side. This viewpoint was first raised by Sazanov and colleagues, who based on the high-resolution cryo-EM structures of Mrp (Multidrug resistant and pH dependent) antiporter [46] and of complex I [47], suggested that only ND5 subunit is well-connected with the periplasmic/inter membrane space (P-side), whereas ND2 and ND4 are not. This in turn raises an intriguing question whether the recruitment of protons from the N side is the only function of ND2 and ND4 antiporter-like subunits?

The higher resolution structures are constantly sought after because they tend to provide accurate position of structural waters due to better interpretation of the cryo-EM densities [25, 48]. MD simulations performed on higher resolution structures are also observed to be more stable [49, 50]. In general, longer simulations with multiple replicates are required to make statistically robust predictions. However, the fixed-charge nature of MD simulations can lead to artificial conformational states upon extensive sampling, especially in case of low-resolution structures. For this reason, atomistic MD simulations were performed on one of the highest resolution complex I structures from *Y. lipolytica* (∼2.1 Å local resolution) [31]. These MD simulations, performed in two different charge states with 3 μs of sampling in each [29], showed no considerable hydration in the P side regions of ND2 and ND4 (Fig. 1B), whereas the putative proton exit route in ND5 was readily formed with water molecules. This data, in agreement with the interpretation from cryo-EM structures, led to a proton pumping proposal where only ND5 acts as the proton releasing route [29] (see also [28]). Importantly, this and the other mechanistic models of only-ND5 pumping are at odds with an earlier report [51], which showed that the truncation of respiratory complex beyond ND2 subunit did not eliminate proton pumping, suggesting that each of the antiporter-like subunits may pick and release protons (see also [52]). In support of this, Beghiah and colleagues [53] recently reconstituted individual antiporter-like subunits in liposomes, and by performing kinetic proton transfer experiments on wild-type and mutant enzymes suggested that each of them are self-contained proton transferring units. Therefore, all in all, the issue that ND2 and ND4 possess proton release routes like ND5, remains unresolved. The discrepancy could stem from the limitations of MD simulations; short time scales (microseconds), limited accuracy of force fields and fixed-charge nature of classical simulations, which may prevent formation of hydrated P-side routes in ND2 and ND4. It also remains possible that cryo-EM models cannot trap the conformational states in which ND2 and ND4 subunits exhibit hydrated P side paths.

A recent high-resolution (2.13 Å) structure of complex I, resolved as part of the *Arabidopsis thaliana* CI:CIII_2_ supercomplex, may have captured such a state [32]. It is the first structure that shows water molecules in the P side region of the ND2 subunit suggesting a possible proton exit route (Fig. 1C). However, as discussed in ref. [29], the path lies deep within the ND2 subunit rather than at the ND2/ND4 interface. Second, it does not link the terminal lysine residue (Lys410, *A. thaliana* complex I numbering) of the ND2 subunit to the P side, as has been proposed based on structural and simulation studies (see above); instead, the path originates from a histidine residue (His236) and progresses towards the P side involving water molecules and polar residues (Fig. 1C). The residues participating in the path are not conserved. The path also coincides with the shift in transmembrane helix 12 (TMH12), which likely creates space for water molecules. Therefore, this structure provides the first structural support for a P-side proton transfer pathway in ND2 subunit of complex I, but its role in proton transfer remains to be established by functional studies.

In this work, inspired by the high-resolution structure of *Arabidopsis* complex I, and building upon our previous MD simulation studies on *Yarrowia* complex [29, 31], we explored the conformational dynamics of the P-side region of ND2 subunit in an intact and a truncated model of the enzyme. We find that an *Arabidopsis* complex I-like conformational change occurs in yeast complex MD simulations, and that the transition occurs much more rapidly in a truncated model constructed to mimic the enzyme studied in earlier biochemical studies [51]. Free energy calculations show that the conformational transition in ND2 subunit occurs much more favourably in the absence of neighboring ND4 subunit with increased hydration in the region. We further explore what triggers the conformational rearrangement, discuss functional significance of this region, and identify residues that can be targeted by site-directed mutagenesis experiments.

## 2. Computational Methods

### 2.1 Unbiased MD simulations

The atomistic MD simulations were based on the high-resolution structure of *Y. lipolytica* complex I (PDB 7O71) and included the protein, cofactors, and structural lipids and water molecules. The protein complex was embedded in a mixed lipid bilayer composed of POPC, POPE, and cardiolipin, solvated with TIP3P water molecules and 100 mM NaCl. The CHARMM36 force field was used for the protein [54], lipids [55], water, ions and cofactors [56–59]. MD simulations were performed following the simulation protocol described by Parey, Lasham et al. [31]. Briefly, the protein-lipid-solvent system was energy-minimized and subsequently equilibrated before the production simulations. Equilibration and production simulations were performed using GROMACS 2022.4 [60] under periodic boundary conditions. During production, the temperature was maintained at 310 K using the Nosé–Hoover thermostat [61, 62] and the pressure at 1 bar using the Parrinello–Rahman barostat [63]. A 2-fs integration timestep was used with the LINCS algorithm [64] for bond constraints. Two different charge states of the protein were simulated, S and P. In state S, all titratable amino acids were modelled in their standard charged states i.e. Asp and Glu were negatively charged and Arg and Lys positively charged. In state P, protonation states were assigned according to the pKa values predicted from PROPKA [65] calculation (see also [31] and Table 1). Two additional setups were created in which the protein structure was truncated (see Table 1) beyond ND2 subunit to reflect on the earlier biochemical data on complex I with distal proton pumping modules deleted [51]. The subcomplex was wrapped in a lipid-solvent environment as before [31] and MD simulations were performed in both charge states S and P (Table 1). An additional *Yarrowia* complex I model system was constructed with the ND2 subunit homology modelled using MODELER [66] based on the structure of *Arabidopsis* complex I subunit ND2. MD simulation trajectory analysis was performed using VMD [67] and MD analysis [68].

**Table 1.** Simulation setups and simulation lengths.

| Complex model | Charge state -<br>Standard (S)<br>Propka (P) | Unbiased simulations -<br>Simulation length x no.<br>of replicas | Biased simulations -<br>Simulation length x no. of<br>replicas |
| --- | --- | --- | --- |
| Full (F) | S | 1 $\mu\text{s}$ x 3 [31] [29] | 100 ns x 3 (AWH) |
| | P | 1 $\mu\text{s}$ x 3 [31] [29] | 150 ns x 3 (AWH) |
|  | S - restrained | - | 180 ns x 5 |
|  | S - homology model | - | 180 ns x 3 |
| Truncated (T) | S | 1 $\mu\text{s}$ x 3 | 100 ns x 3 (AWH) |
|  | P | 300 ns x 3 | 100 ns x 3 (AWH) |

### 2.2 Restrained simulations and free energy calculations

We performed free energy calculations using the AWH (Accelerated Weight Histogram) method [69], starting from the final snapshot of the NPT equilibration step [31]. AWH simulations were performed for four states (F/S, F/P, T/S, and T/P, where T – truncated and F – full, see Table 1). The simulations were performed using the multiple-walker method implemented in GROMACS (awh-multi) using four walkers. The Cα–Cα distance between Asn385 and Ser308 (*Y. lipolytica* complex I numbering) was used as the reaction coordinate, with a force constant of 12 kcal mol⁻¹ Å⁻², and the distance was varied from 4 Å to 8 Å. Each simulation replica was continued until convergence was reached (see Supplementary Information, Fig. S3), and only the final 50–100 ns of each simulation were used for analysis.

Inspired by the proximity of sidechains of Lys410 (ND2) and Glu147 (ND4) from TMHs 12 and 5, respectively, in *Arabidopsis* complex I structure (PDB 8BEF), we modelled the equivalent Lys383 and Glu142 sidechains in hydrogen bonding distance in the *Yarrowia* complex I by applying a harmonic restraint (∼36 kcal mol⁻¹ Å⁻²) between the Nζ (Lys) and Cδ (Glu) atoms with a biased distance of 3.2 Å and performed MD simulations (see Table 1).

MD simulations of full *Y. lipolytica* complex I with its ND2 subunit homology modelled based on plant complex conformation were performed with Cα atoms of the modelled subunit restrained to their positions by harmonic restraints of ∼48 kcal mol⁻¹ Å⁻².

## 3. Results

### 3.1 MD simulations of Y. lipolytica complex I capture unique conformational changes

To probe the conformational dynamics of the region that may be responsible for proton exit in the ND2 subunit, we scrutinized the long (in total 6 μs) MD simulation trajectories of complex I from *Y. lipolytica* in its two charge states S and P. Throughout the simulations, the overall protein structure remains stable (Fig. S1), and no major global conformational changes are observed. However, noticeable local rearrangements occur at the residue level. Among these, a salient transition involves two polar residues Asn385 and Ser308 near the P side region of the ND2 subunit (Fig. 2A). In MD simulations, the distance between the backbone Cα atoms of these two residues shifts from a shorter *d*_S_ ∼ 5.2 Å to a longer *d*_L_ ∼ 6.5 Å (Fig. 2B and S2). Once the conformation associated with longer distance is reached, it remains stable for the rest of the simulation time (Fig. S2). This stability is achieved in part due to the rotation of the sidechain of Asn385, which establishes a hydrogen bond with the sidechain of Ser308 (Fig. 2C). The conformational dynamics of the Asn-Ser pair is found to be dependent on the charge state of the protein. In the P state simulations, the Cα-Cα distance rapidly increases with the transition occurring within the first 400 ns in every single replica (Fig. S2). In contrast in the S state simulations the transition is much slower. Only in one simulation replica, the distance increases to *d*_L_ ∼ 6.5 Å within 400 ns, while the other replicates show either no conformational change or occurring much later in the simulation.

**Fig. 2.**
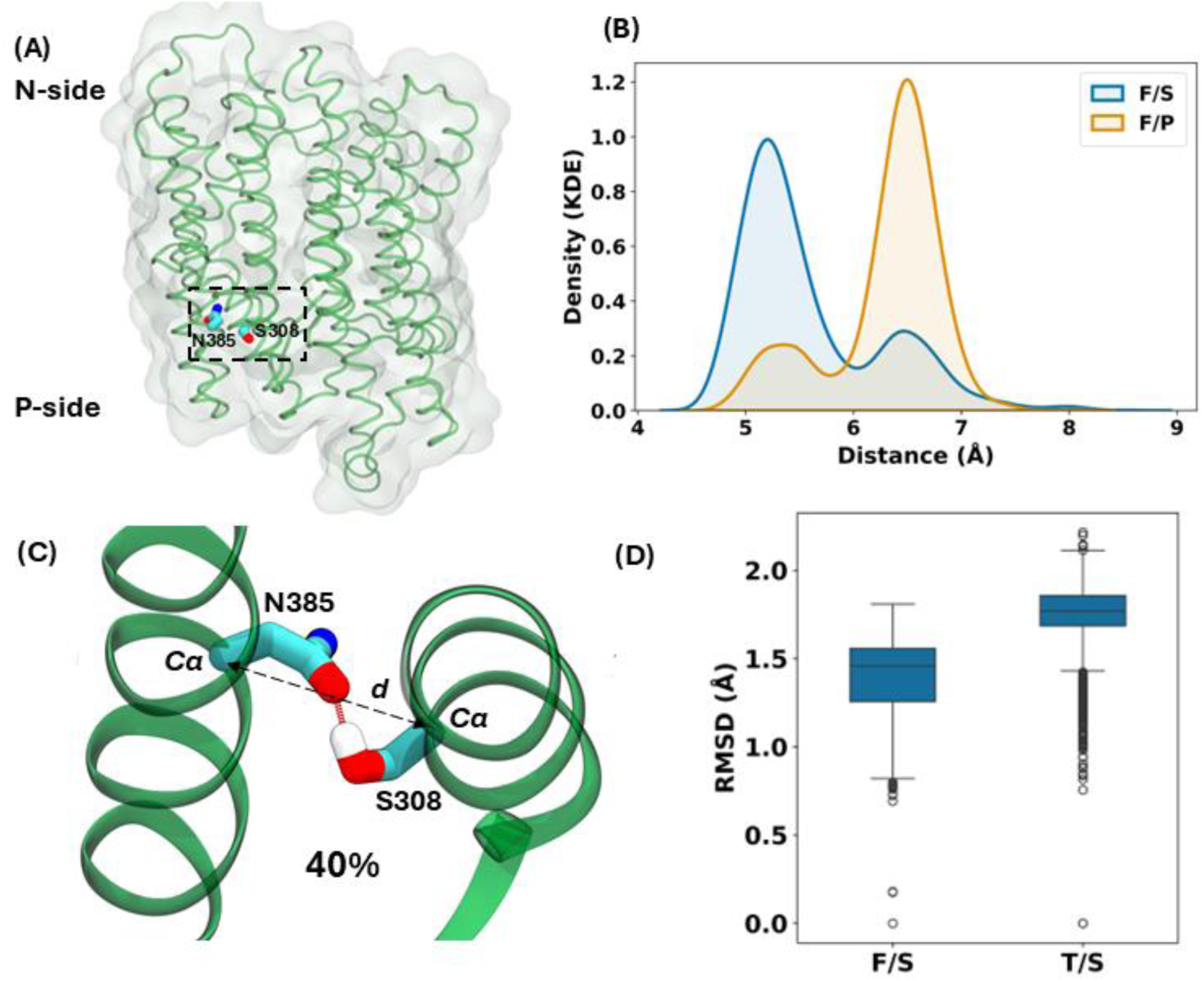
Conformational dynamics in the ND2 subunit of complex I. (A) ND2 subunit is displayed with ribbons and the region undergoing conformational transition (Asn385 and Ser308) is shown in licorice representation (dotted box). The sidechains of two residues are shown as in the cryo-EM structure (PDB 7O71). (B) Kernel Density Estimate (KDE) of the Asn385-Ser308 Cα-Cα distance (defined in panel C) from MD simulations of full complex I in S and P charge states. (C) Rotation of asparagine sidechain established a hydrogen bond between the sidechains of Asn385 and Ser308 with an occupancy of ca. 40% in unbiased F/P simulations. The hydrogen bond between rotated Asn385 and Ser308 is shown with a red dotted line. (D) RMSD of the Cα backbone atoms of the ND2 subunit in S state calculated with respect to the first frame of each production trajectory. S – standard and P – propka based charged states, whereas F and T represent full and truncated complex I (see methods).

The conformational dynamics of Asn-Ser pair is interesting because the longer distance *d*_L_ ∼ 6.5 Å resembles the Cα-Cα distance of Tyr-Ser pair in the ND2 subunit of complex I from *Arabidopsis thaliana* (∼6.6 Å, Fig. 2B). The two residues Tyr412 and Ser344 (*Arabidopsis* complex I numbering) occupy the same position as Asn385 and Ser308 in *Yarrowia* complex I and line the region around structurally resolved water molecules in plant complex I (Fig. 1C). In *Y. lipolytica* complex I simulations no stable water molecules are found in this region in the timescales of the simulation, even though the Asn-Ser associated conformational change creates enough space to accommodate water molecules (see more below).

### 3.2 Long time scale dynamics of truncated complex I

Previous biochemical experiments on *Y. lipolytica* complex I [51] suggested that truncated complex I retains sub-stoichiometric proton pumping activity despite lacking the distal proton pumping modules (see Introduction). In contrast, data on bacterial complex from *E. coli* showed that genetic deletion of ND5 and ND4 subunits results in an unstable complex, which shows no quinone reductase activity [52] (see also [70, 71]). In the background of this conflicting data, we envisaged a scenario in which truncated complex I creates an artificial proton pumping route in the membrane exposed ND2 subunit, which is otherwise blocked in an intact enzyme. To test this hypothesis, we carried out MD simulations of a truncated complex I in which all subunits beyond ND2 were removed *in-silico* (see methods and Table 1) and probed the dynamics of Cα-Cα distance and hydration in the region as a proxy for putative proton transfer route towards the P-side. We find that the transition from shorter to longer Cα-Cα distance occurs rapidly in all the simulation trajectories and in both states (T/P and T/S), with the conformational change occurring in less than 50 ns (Figs. 3A and S2). The density plots further reveal that none of the truncated model systems explore short distance (*d*_S_) population, in clear contrast to the full-length structure in the S state, where the shorter Cα-Cα distance was much more populated (Fig. 2). Overall, these data indicate that truncation of complex I favors the ND2 associated conformational change. Moreover, unlike the F/S and F/P state simulations of intact enzyme, the truncated system displays hydration in the region near Asn385 and Ser308 (see Fig. 3B), similar to what has been observed in the ND2 subunit of complex I from *A. thaliana* (Fig. 1D). Importantly, the hydration in this region is largely buried in the ND2 subunit and is not at the interface with the lipid membrane (see Discussion).

**Fig. 3.**
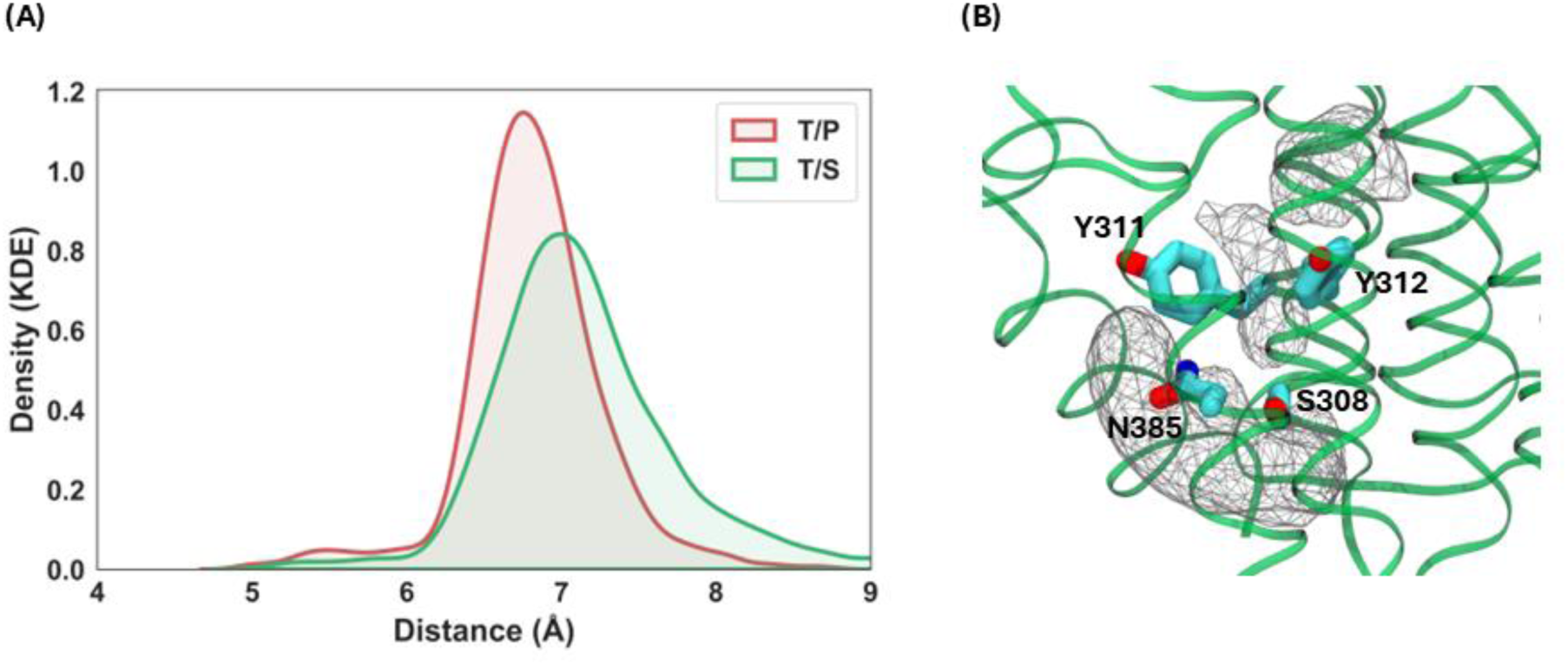
Dynamics of the subcomplex. (A) Density distribution of the Cα-Cα distance (Asn385 and Ser308 of ND2 subunit) in truncated (T) models of complex I in two charge states P and S. (B) Hydration in the P side region of ND2 subunit shown as a water occupancy map (displayed at 0.2 isovalue) from MD simulation of truncated complex I in S state and selected amino acid residues.

### 3.3 Energetic estimates on the ND2 conformational change

Given that the ND2-associated conformational change occurs relatively fast in the MD simulations of truncated complex I, we estimated the energetic stabilization provided by the protein in full complex by performing AWH [69] free energy simulations (see methods). To this end, we selected a simple collective variable (Cα-Cα Ser-Asn distance) and performed multiple independent AWH simulations, which all converged within 100-150 ns time frame (Fig. S3). We find that full complexes in P and S charge states prefer the short-distance (*d*_S_ ∼ 5.2 Å) regime (Fig. 4A), whereas a truncated complex I is equally likely to be found with Asn-Ser pair Cα atoms separated by ∼6.5 Å (*d*_L_) (Fig. 4B). The state with longer Cα-Cα distance (*d*_L_) is ca. 5 kcal/mol higher in energy in the full complex. In contrast, both the short- and long-distance states are equally likely in truncated complex I (Fig. 4). In other words, truncation abolishes the free energy difference between the short- and long-distance states consistent with the higher flexibility of the truncated system observed in unbiased simulations (Fig. 2D). The flexibility-induced conformational change and the resulting hydration (Fig. 3) is in part due to the membrane exposure of the ND2 subunit. In truncated complex I, ND2 resembles the membrane-facing terminal antiporter-like subunit ND5 that displays a strong hydration pattern towards the P side of the membrane (see also [31, 45]). In contrast, in intact complex I, ND2 is in contact with the ND4 subunit, which can mechanically restrain its conformational dynamics.

**Fig. 4.**
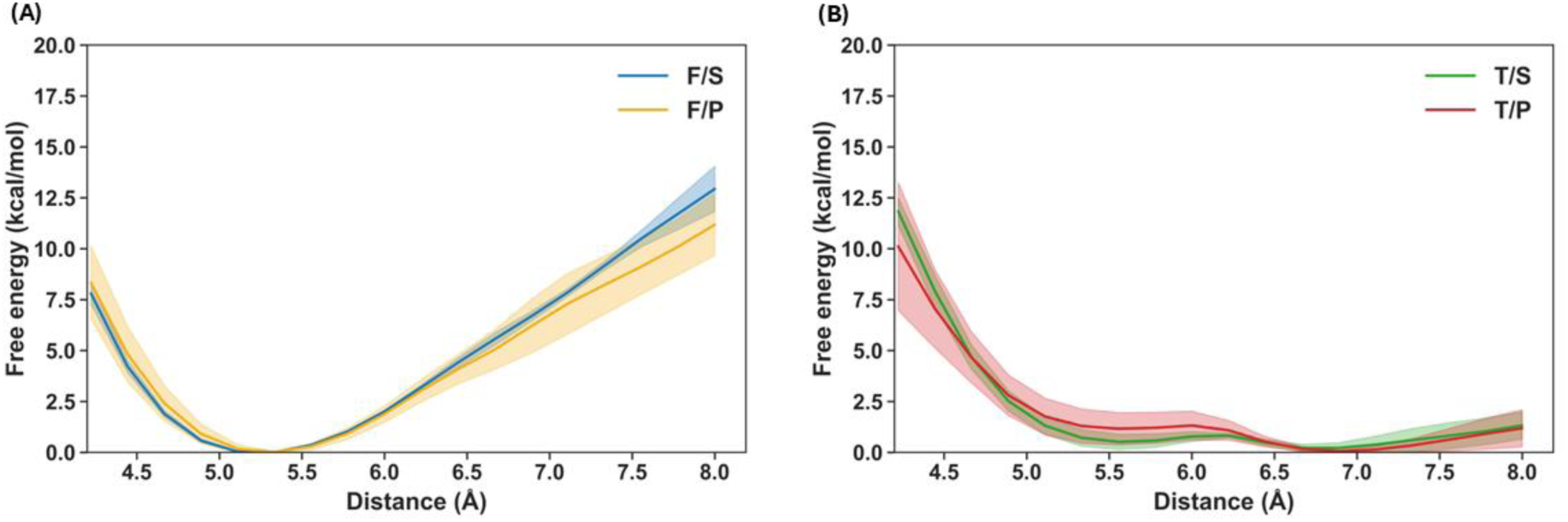
Energetics associated with the change in Asn385-Ser308 Cα-Cα distance. (A) Full complex and (B) truncated complex I in P and S states. In panels (A) and (B), the y-axis displays the free energy (in kcal/mol) for the Asn-Ser Cα-Cα distance in Å (x-axis). The traces represent the average of three simulation replicas, and the light-shaded areas denote the standard error of the mean (SEM).

### 3.4 Possible triggering mechanisms for ND2 conformational transition and correlated changes

The above results suggest that truncation of complex I triggers the conformational transition in the P side region of the ND2 subunit. However, it remains unclear what structural features would govern this conformational change in an intact complex I. We noticed that in plant complex I structure, the lysine residue (Lys410) from the ND2 subunit points towards the acidic residue (Glu147) of the ND4 subunit (Fig. 5A). The two residues establish a water-mediated interaction forming a polar bridge within the hydrophobic surroundings of two antiporter-like subunits. In contrast, the two residues are 12-13 Å apart in *Yarrowia* complex I (Fig. 5A). By hypothesizing that proximity of lysine and glutamate can play a role in triggering the conformational change in ND2 subunit, we performed MD simulations in which the distance between Lys383 and Glu142 (*Y. lipolytica* numbering) was harmonically restrained to 3.2 Å (see methods and Table 1, F/S - restrained), thereby mimicking the arrangement observed in *Arabidopsis* complex I. We observed that the Asn385 – Ser308 distance increases up to ∼7 Å in MD simulations with modelled interaction between Lys383 and Glu142 (Fig. 5B). Even though the conformational change occurs rarely (ca. two out of five simulation replicas), analysis of trajectories shows transient diffusion of water molecules in the region when Cα-Cα increases to larger values (Fig. 5C) similar to what is observed in truncated complex I simulations.

**Fig. 5.**
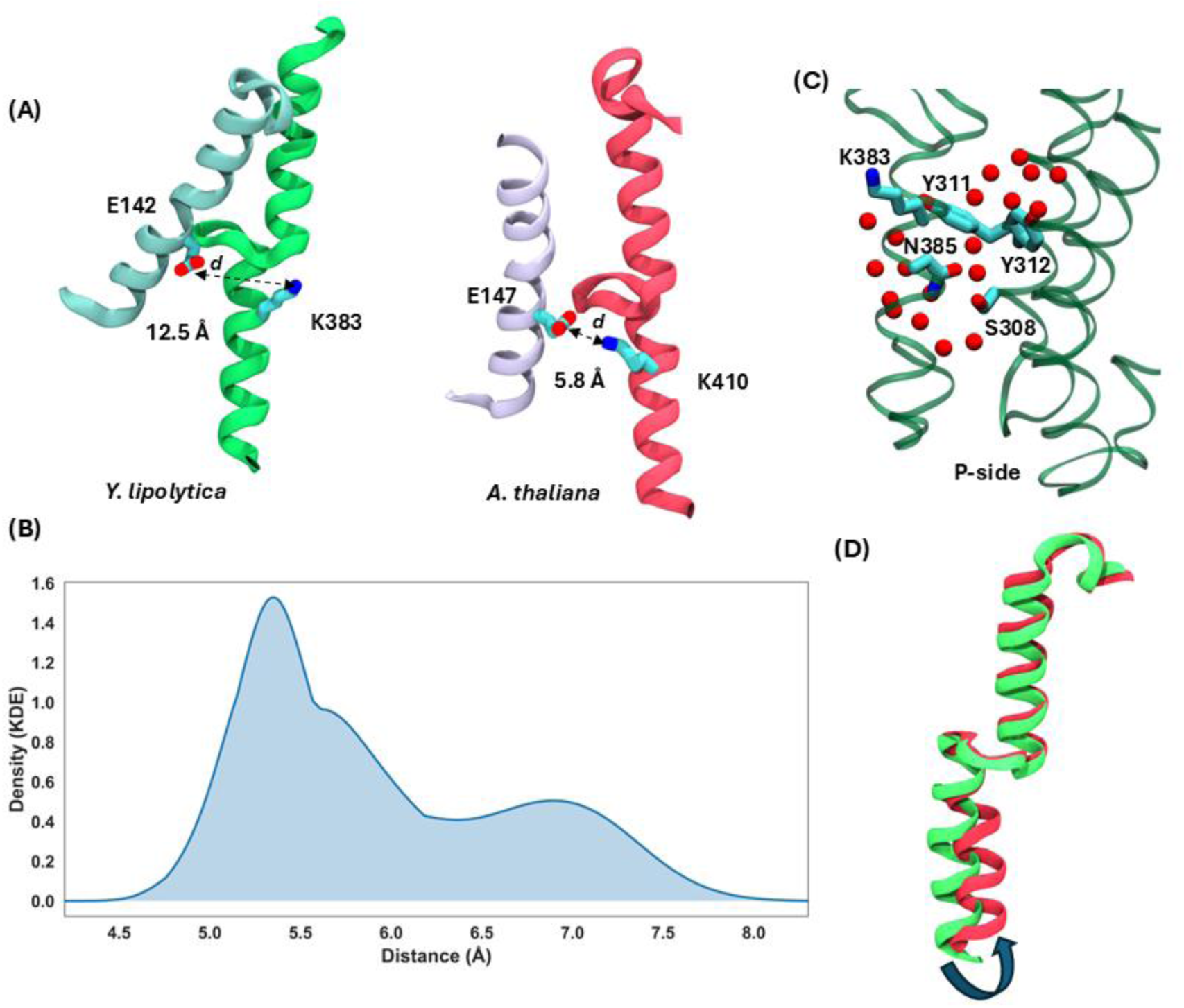
Structural features triggering Asn-Ser pair dynamics. (A) Conserved lysine residue from TMH12 of ND2 subunit and acidic residue from TMH5 of ND4 subunit shown from *Y. lipolytica* and *A. thaliana* structures. (B) KDE plot of Asn385 – Ser308 Cα-Cα distance from MD simulations where Lys383 (ND2) and Glu142 (ND4) are modeled close to each other (see methods). (C) Simulation snapshot showing water molecules (red spheres) enter the region upon increase in Asn385 – Ser308 Cα-Cα distance in MD simulations with sidechains of Lys383 and Glu142 harmonically restrained towards each other. (D) TMH12 of ND2 subunit from *Yarrowia* and *Arabidopsis* complexes. Arrow shows displacement in part of the helix.

An additional noticeable structural difference between the two structures is that part of the TMH12 is shifted in plant complex I compared to the yeast structure (Fig. 5D). The reason for this difference is unclear but could in part be related to the bending of the membrane domain of complex I, which in this structure is part of a CI:CIII_2_ supercomplex or due to the presence of a detergent molecule present nearby. By envisaging that the shift of TMH12 could trigger Asn-Ser pair conformational change, we simulated a full complex I model with ND2 subunit homology modelled based on the same subunit from plant complex I (Table 1, setup F/S - homology model). Due to the re-conversion of the homology modelled ND2 to original yeast-like conformation, we applied harmonic restraints to the Cα atoms of the ND2 subunit. The plant complex I-like conformation with an average Asn-Ser Cα-Cα distance of ∼6.2 Å is maintained in the restrained simulation but hydration of the region is not observed, likely due to the limited time scales (∼500 ns in total) and restricted conformational dynamics of the system.

We further examined whether a pattern like Asn-Ser conformational change can be observed in other residues of the ND2 subunit. Through contact map analysis, we identified several pair of residues that display increased Cα-Cα distance in a truncated (T/S) but not in intact (F/S) complex I simulation (Table S1). For instance, in the proximity of Ser-Asn pair, the Cα-Cα distance between Tyr312 and Gly382 is found to show a marked increase, suggesting concerted changes can also take place elsewhere in the ND2 subunit (Table S1, see Discussion).

## 4. Discussion

The mechanism of proton pumping by respiratory complex I is one of the key unresolved problems in molecular bioenergetics. The membrane-bound subunits ND2, ND4 and ND5 are responsible for proton pumping, which occurs in response to the exergonic reduction of quinone by NADH. Despite extensive mutagenesis data on complex I, and its 3D structures and computer simulations, the molecular description of proton pumping by ND2/4/5 subunits remain unresolved and much debated (see Introduction). The general assumption is that each of the three antiporter-like subunits has a proton uptake pathway that connects the N side of the membrane to the middle of the protein subunit. There is an emerging agreement in the field that a proton exit pathway is present in terminal ND5 subunit. However, the question remains as to whether ND2 and ND4 subunits also contain similar proton release routes or if they are solely involved in proton uptake. If one assumes the only proton release route is through ND5, one can expect a subcomplex without ND5 subunit to be devoid of proton pumping. But this is not the case; a *Yarrowia lipolytica* complex I, in which distal subunits ND5 and ND4 have been removed genetically, pumps protons, albeit at a lower H^+^/e^-^ stoichiometry [51]. One can hypothesize a scenario in which truncation of ND4/ND5 leads to the creation of an artificial proton exit route in the ND2 subunit, which otherwise does not exist in an intact enzyme (see also [31]). To test this possibility, we created a truncated model of *Y. lipolytica* complex I, and compared the conformational dynamics of its ND2 subunit with the same in an intact complex I. The loss of neighboring ND4 subunit and exposure of the ND2 subunit to lipid bilayer results in a conformational change that supports hydration in the region and formation of a pathway that may participate in proton transfer to the P side. Free energy estimates suggest loss of neighboring protein subunit reduces the energetic costs for this conformational change, which involves rotation of the sidechain of Asn385 and establishment of a hydrogen bond with Ser308 from the same subunit ND2. This causes the backbone Cα atoms of the two residues to move apart by 1-2 Å, creating extra space that fills with water molecules depending on the charge state of the protein. Interestingly, in a recently solved 2.13 Å cryo-EM model of plant complex I (PDB 8BEF, [32]), the same region is found to hold structurally-observed water molecules. We find the Cα atoms of two residues Tyr412 and Ser344 (same position as Asn385 and Ser308 in the ND2 sequence) are separated by a similar distance as observed in MD simulations of *Yarrowia* complex I. In other words, our MD simulations are able to capture a conformational transition and hydration in yeast complex I, similar to what is observed structurally in plant complex I despite variation in the local amino acid environment. However, the question remains whether the observed conformational changes represent formation of proton release route to the P-side or are they just incidental structural transitions to accommodate functional changes in other parts of the protein. When high-resolution structures of other complexes are analyzed, a similar variation in distance of Cα atoms is observed (Table 2), even though the residues involved in these positions are not conserved among complex I from different species (see also [29]). Therefore, if the region is indeed responsible for proton exit in the ND2 subunit, it is somewhat hard to fathom why the amino acid composition is not conserved and the residues are even replaced by non-polar counterparts. Alternatively, if this is not a proton pumping route, what purpose do these conformational transitions serve? In that case, these may represent conformational response to functionally relevant transitions elsewhere in the protein. The conformation-coupled hydration could assist in enzyme catalysis in an indirect manner; for instance, hydrated regions in proteins are known to act as dielectric wells [72, 73], where they respond to charge changes, but do not transport protons.

**Table 2.** Analysis of Cα-Cα distance between residues from ND2 subunit in selected structures. *Yl – Yarrowia lipolytica; At – Arabidopsis thaliana; Bt – Bos taurus; Ec – Escherichia coli; Ct – Chaetomium thermophilum; Mm – Mus musculus; Ss – Sus scrofa; Oa – Ovis aries; Te – Thermosynechococcus elongatus; Tv – Thermosynechococcus vestitus;

| PDB ID | ~ Resolution (Å) | Organism* | Residue 1 | Residue 2 | ~ Ca-Ca Distance (Å) |
| --- | --- | --- | --- | --- | --- |
| 7O71 | 2.4 [31] | *Yl* | Asn385 | Ser308 | 5.2 |
| 8BPX | 2.09 [32] | *At* | Tyr412 | Ser344 | 6.6 |
| 7QSM | 2.3 [74] | *Bt* | Met265 | Met200 | 6.9 |
| 7Z7V | 2.29 [25] | *Ec* | Tyr397 | Ala325 | 5.1 |
| 7ZMB | 2.75 [75] | *Ct* | Met453 | Ala365 | 5.2 |
| 6ZTQ | 3.0 [76] | *Mm* | Ile265 | Leu200 | 7.7 |
| 8OM1 | 2.39 [77] | *Mm* | Ile265 | Leu200 | 7.5 |
| 7VYS | 2.5 [23] | *Ss* | Met265 | Met200 | 7.1 |
| 6ZKA | 2.5 [47] | *Oa* | Met265 | Met200 | 6.7 |
| 6HUM | 3.34 [78] | *Te* | Tyr397 | Ser328 | 6.0 |
| 6KHJ | 3.0 [79] | *Tv* | Tyr397 | Ser328 | 5.4 |
| 6TJV | 3.2 [80] | *Te* | Tyr397 | Ser328 | 5.5 |

To probe the functional importance of the region, it would be reasonable to exchange the polar residues with the non-polar ones and assess the quinone reductase activity and proton pumping by complex I. To the best of our knowledge, residues in this region of complex I have not been studied with site-directed mutagenesis studies. Alternatively, the conformational dynamics of the region can be restricted by introducing cysteine residues and an artificial covalent linkage and evaluating enzymatic activity. Conformational locking of a conserved loop in complex I by a cysteine-based covalent linkage has been shown to affect proton pumping but not the reduction of Q [81, 82].

The above-described hydrated region of ND2 subunit in plant and yeast complexes is well buried within the protein matrix. However, an alternative path towards the P side could be expected at the interface of ND2 and ND4, similar to the proton exit route in membrane-facing ND5 subunit [29, 31]. Interestingly, an array of water molecules is observed in *Arabidopsis* complex I structure approaching the antiporter-antiporter interface in part due to the space created by shift in TMH12 (Fig. 5) and surrounding helices. The displacement of TMH12 causes a small change in the position of hydrophobic residue Phe411 (plant complex I numbering, Leu384 in *Yarrowia* complex I), which may cause the entry of water molecules in the region. The residue position is occupied by a hydrophobic residue in ND2 and ND4 subunits but is replaced by a conserved acidic residue in ND5 subunit (Asp397 in *Y. lipolytica* complex I). The latter residue has been suggested to form a proton release group towards the P side of the membrane based on QM/MM MD simulations of bacterial complex I [45]. When hydrophobic residues of yeast ND2 and ND4 subunits (Leu384 and Phe396, respectively) are replaced *in silico* to mimic the acidic site of ND5 (an aspartate), rapid hydration is seen in the interfacial region between ND5 and ND4, and ND4 and ND2 subunits, also linking the central hydrophilic axis to the P-side of the membrane [31]. This suggests that hydrophobic blockages in ND2 and ND4 subunits insulate the central hydrophilic axis of complex I and also the subunit-subunit interface, but upon structural [32] or chemical perturbation [31] may create hydrated pathways.

If ND2 and ND4 subunits do not carry proton releasing routes towards the P side, this would require several mechanistic models of proton pumping by complex I to be revised. The photosynthetic NDH-1 complexes provide an interesting perspective in this regard. In NDH-1 complexes of the type NDH-1_4_ (NDH-1MS’), the residues responsible for P side proton exit are not conserved in the ND5 homologue NdhF4 [83], suggesting this release route may be blocked. If we assume ND2 and ND4 subunits do not transport protons to the P side of the membrane (cf. [84]), it remains unclear what happens to the energy released upon plastoquinone (PQ) reduction from ferredoxin (∼ -24 kcal/mol) in NDH-1_4_ complex. Part of it may be used to pump protons across the membrane as has been suggested recently [84]. This could occur via the fourth proton pumping route that has been proposed based on structural and simulation studies on canonical complex I [24, 42, 43] and may not involve the three antiporter-like subunits. It can also be speculated that there is a direct coupling between the PQ reduction and CO_2_ hydration reaction catalyzed by the enzyme; in this case the substrate protons required for two-electron reduction of PQ could in part be provided by the protons released upon bicarbonate formation. The transfer of protons, instead of occurring through the cytoplasmic space, may occur through the central hydrophilic axis, which in the recent high-resolution structure of NDH-1MS’ [83] connects the CupB domain to the PQ reduction site. Interestingly, coupling of CO_2_ hydration reaction with the proton pumping function of complex I has been suggested based on plant complex I structure; in *Arabidopsis* complex a carbonic anhydrase domain (γCA) is anchored to the cytoplasmic side of the ND2 subunit and may assist in funneling protons for redox-coupled proton pumping [85].

## 5. Concluding remarks

In this work, by performing atomistic MD simulations of high-resolution cryo-EM structure of complex I from *Yarrowia lipolytica*, we have identified a unique conformational change in the P-side region of its ND2 antiporter-like subunit. Among others, the conformational transition involves residues Asn385 and Ser308 from the ND2 subunit, where they establish a hydrogen bond between their sidechains leading to the shift of their Cα atoms. Free energy calculations show that this transition occurs much more favourably in the absence of neighboring ND4 subunit, which also leads to the entry of water molecules in the region near Asn-Ser pair. Given the weak sequence conservation of the region, it may not represent a proton release route of the ND2 subunit. Instead, the observed conformational change may reflect a secondary response to catalytic transitions elsewhere in the protein.

## Acknowledgements

VS acknowledges research funding from the Sigrid Jusélius Foundation, the Jane and Aatos Erkko Foundation, the University of Helsinki and the Research Council of Finland. The supercomputing resources from Center for Scientific Computing (CSC), Finland are gratefully acknowledged. VS is thankful to Volker Zickerman, Jan Schuller, Marc Nowaczyk and Yat Kei Lo for helpful discussions on mitochondrial complex I and photosynthetic NDH-1.

## Supplementary Information

**Fig S1.**
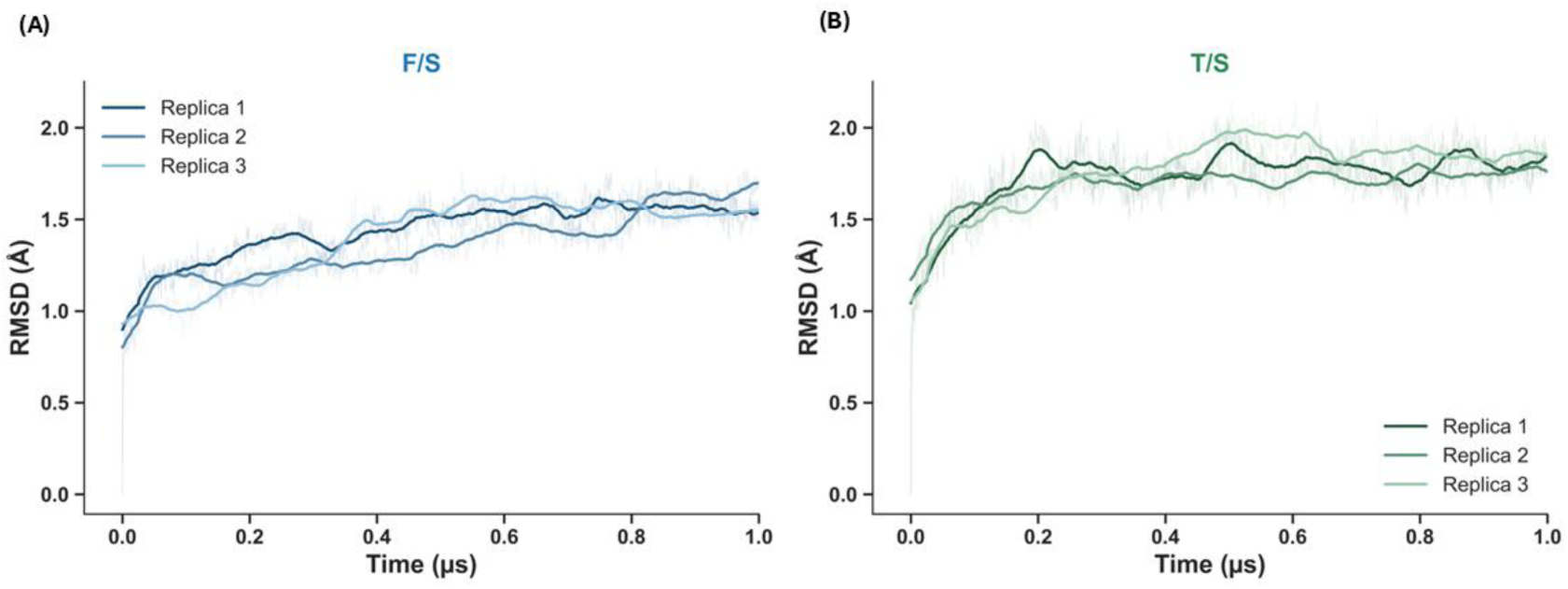
Time series representation of root mean square deviation (RMSD) of backbone atoms of ND2 subunit in F/S (A) and T/S (B) systems, calculated with respect to the first frame of each production trajectory. Three replicas are displayed.

**Fig. S2.**
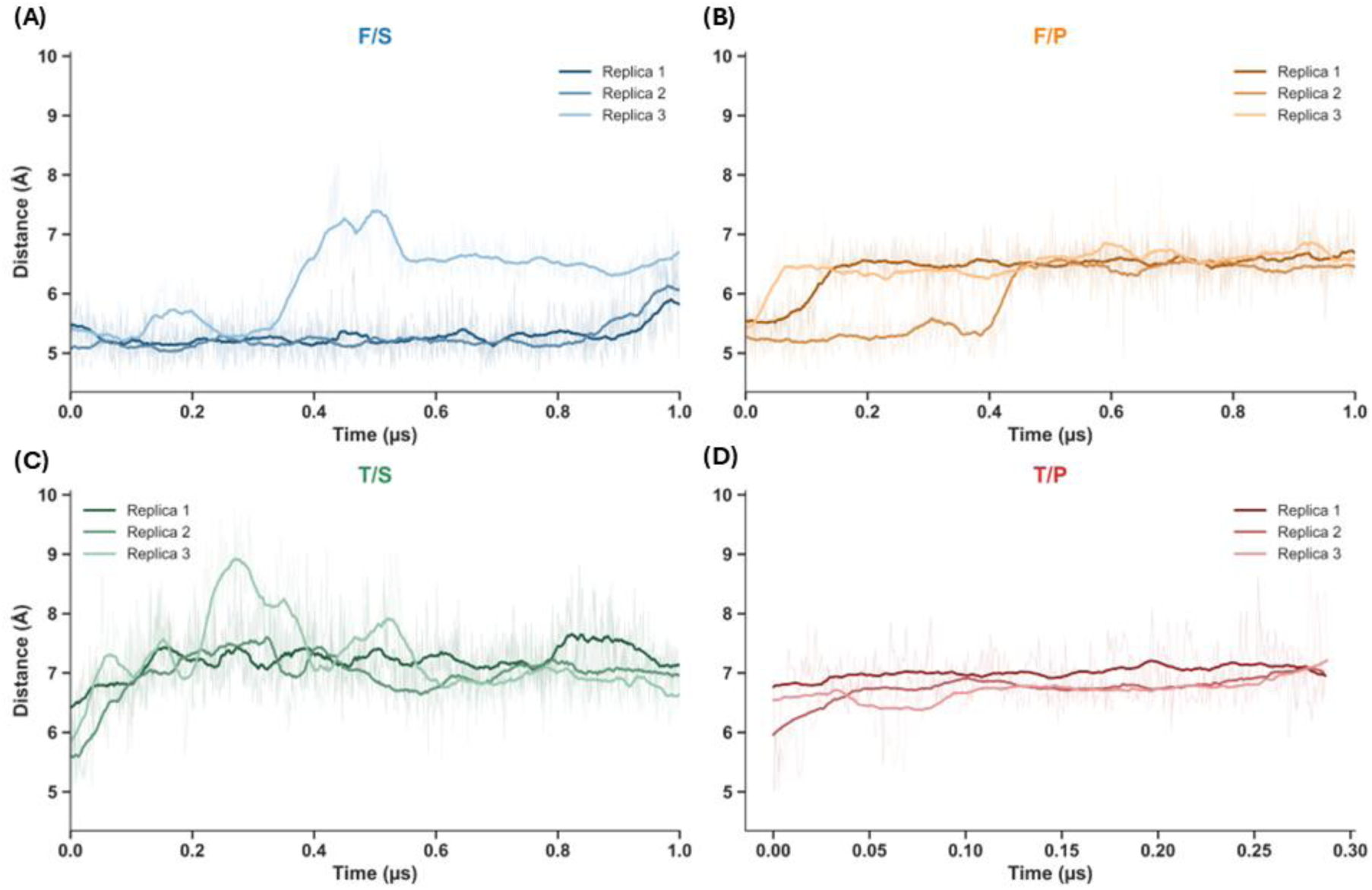
Time series representation of the Cα–Cα distance between the Ser-Asn pair from F/S (A), F/P (B), T/S (C), and T/P (D) simulation setups.

**Fig. S3.**
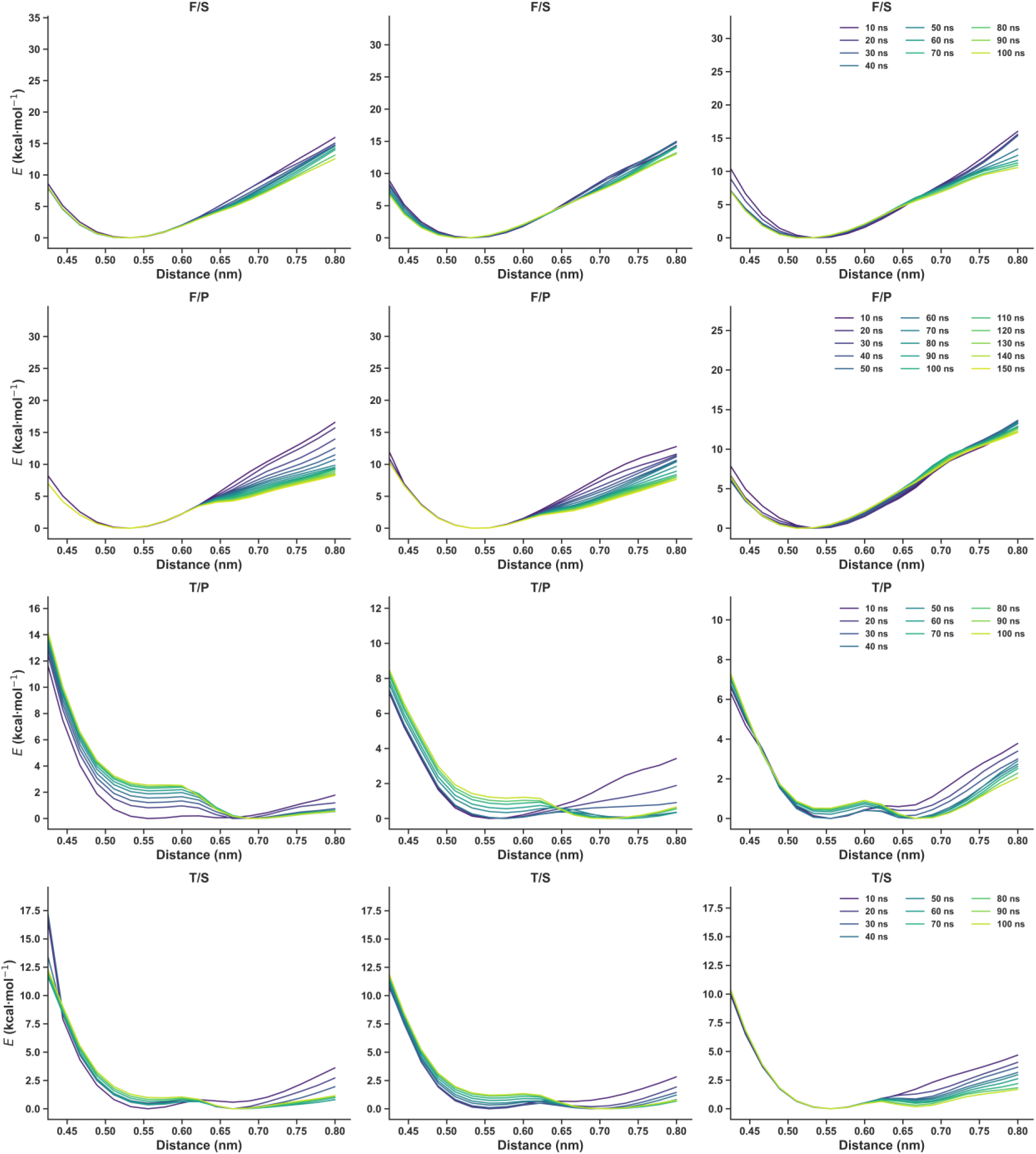
Convergence of the free-energy profiles obtained from AWH simulations for F/S, T/S, F/P, and T/P model systems (three replicas each). The profiles are shown as a function of simulation time to assess the convergence.

**Table S1.** Change of contacts between residue pairs from the ND2 subunit in intact and truncated model systems. Contacts were computed over all simulation frames (F/S and T/S states) with a distance cutoff of 5.3 Å corresponding to the shorter Ser-Asn Cα-Cα distance (*d*_S_, also based on the minima observed in AWH simulations). Δ = contacts (T/S) – contacts (F/S); only pairs with |Δ| ≥ 0.30 are shown. Negative values indicate contacts lost in the truncated system, positive values indicate contacts gained.

| Residue i | Residue j | $\Delta$ |
| --- | --- | --- |
| Val37 | Ser40 | -0.44 |
| Asn48 | Tyr467 | -0.39 |
| Asn49 | Tyr467 | -0.49 |
| Phe159 | Gly162 | -0.51 |
| Asn193 | Asp196 | -0.41 |
| Ser308 | Asn385 | -0.49 |
| Tyr312 | Gly382 | -0.68 |
| Asn338 | Pro343 | -0.33 |
| Lys351 | Val419 | -0.32 |
| Gly352 | Ser420 | -0.46 |
| Gly352 | Gln422 | -0.39 |
| Leu353 | Ser420 | -0.32 |
| His355 | Ile427 | -0.68 |
| Asp356 | Leu428 | -0.79 |
| Tyr359 | Val361 | -0.43 |
| Gly373 | Gly379 | -0.31 |
| Ile400 | Ile403 | -0.31 |
| Asn457 | Ile460 | -0.37 |
| Leu42 | Leu44 | +0.36 |
| Thr95 | Glu98 | +0.30 |
| Tyr148 | Ser151 | +0.35 |
| Leu156 | Phe159 | +0.46 |
| Leu197 | Ile200 | +0.39 |
| Glu226 | Lys282 | +0.37 |
| Ser272 | Gly275 | +0.32 |
| Val273 | Leu276 | +0.49 |
| Leu276 | Tyr412 | +0.32 |
| Tyr295 | Leu298 | +0.42 |
| Met296 | Leu299 | +0.30 |
| Asn305 | Ser308 | +0.45 |
| Gly352 | Asp423 | +0.57 |
| Gly352 | Lys424 | +0.57 |
| Leu377 | Gly453 | +0.57 |
| Leu378 | Gly453 | +0.58 |
| Leu387 | Ile390 | +0.53 |
| Met388 | Leu391 | +0.47 |
| Tyr413 | Leu416 | +0.31 |
| Ile450 | Gly453 | +0.43 |
| Gly453 | Tyr456 | +0.53 |
| Ile455 | Ser458 | +0.39 |
| Tyr456 | Ser458 | +0.31 |

